# Dopamine dips during unrewarded actions promote punishment-resistant reward seeking

**DOI:** 10.64898/2026.08.01.742224

**Authors:** Nkatha Mwenda, Jacob A. Nadel, Tong Shen, Jillian L. Seiler, Talia N. Lerner

## Abstract

Punishment-resistant reward-seeking, a hallmark of addiction, is less prominent in females than males. We found that chronic estradiol manipulations increased punishment resistance in female mice without increasing dopamine peaks on rewarded actions as expected, instead exaggerating dopamine dips on unrewarded actions; optogenetically mimicking these dips accelerated punishment resistance in females and males. These results suggest dopamine dips suppress learning from unrewarded actions, consistent with policy-based accounts of dopamine function.

## MAIN

In humans and rodents, females are, on average, less impulsive, less sensation-seeking, and less punishment-resistant than males ^1–7^. What creates these sex differences in behavior? Previous studies suggest a role for the sex hormone estradiol ^8–12^, which can modulate dopamine signaling ^13–20^.

In our previous work, we showed that the amplitude of phasic dopamine peaks in the dorsomedial striatum (DMS) on rewarded actions during random interval reinforcement training (RI60) predicted the development of punishment-resistant reward seeking, a behavioral hallmark of addiction ^21^. Further, optogenetic stimulation of DMS dopamine peaks during rewarded actions in RI60 training was sufficient to increase punishment-resistant reward-seeking, while optogenetic inhibition of these peaks prevented it ^7^. Importantly, although the optogenetic effects were slightly stronger in male mice, they were effective in both males and females, implicating the DMS dopamine system in this behavior across the sexes.

Here, we investigated whether sex differences in punishment-resistant reward-seeking arise because of estradiol-dependent modulation of DMS dopamine signals in female mice. We used fiber photometry and the fluorescent dopamine sensor GRAB-gDA3m to record DMS dopamine signals during RI60 training in three groups of female mice (Fig. 1a-b, S1a-b). We compared normally cycling females with non-cycling ovariectomized (OVX) females, with or without 17β-estradiol replacement. All mice underwent OVX or sham surgery and subcutaneous implantation of slow-release pellets for placebo or hormone treatment. The first group (Sham), received a sham surgery and a placebo pellet. The second group (OVX-P) received OVX surgery and a placebo pellet to mimic a chronically low estradiol state similar to diestrus. The third group (OVX-E) received OVX surgery and a 90-day slow-release 17β-estradiol pellet to mimic a high estradiol state similar to proestrus (Fig. 1b, S1c). After recovery from surgery, mice received RI60 training with early and late shock probes to test for punishment resistance, as in our previous study (Fig. 1c) ^7^. Behavior in the RI60 task was indistinguishable amongst the groups (Fig. S2a-d).

**Figure 1.**
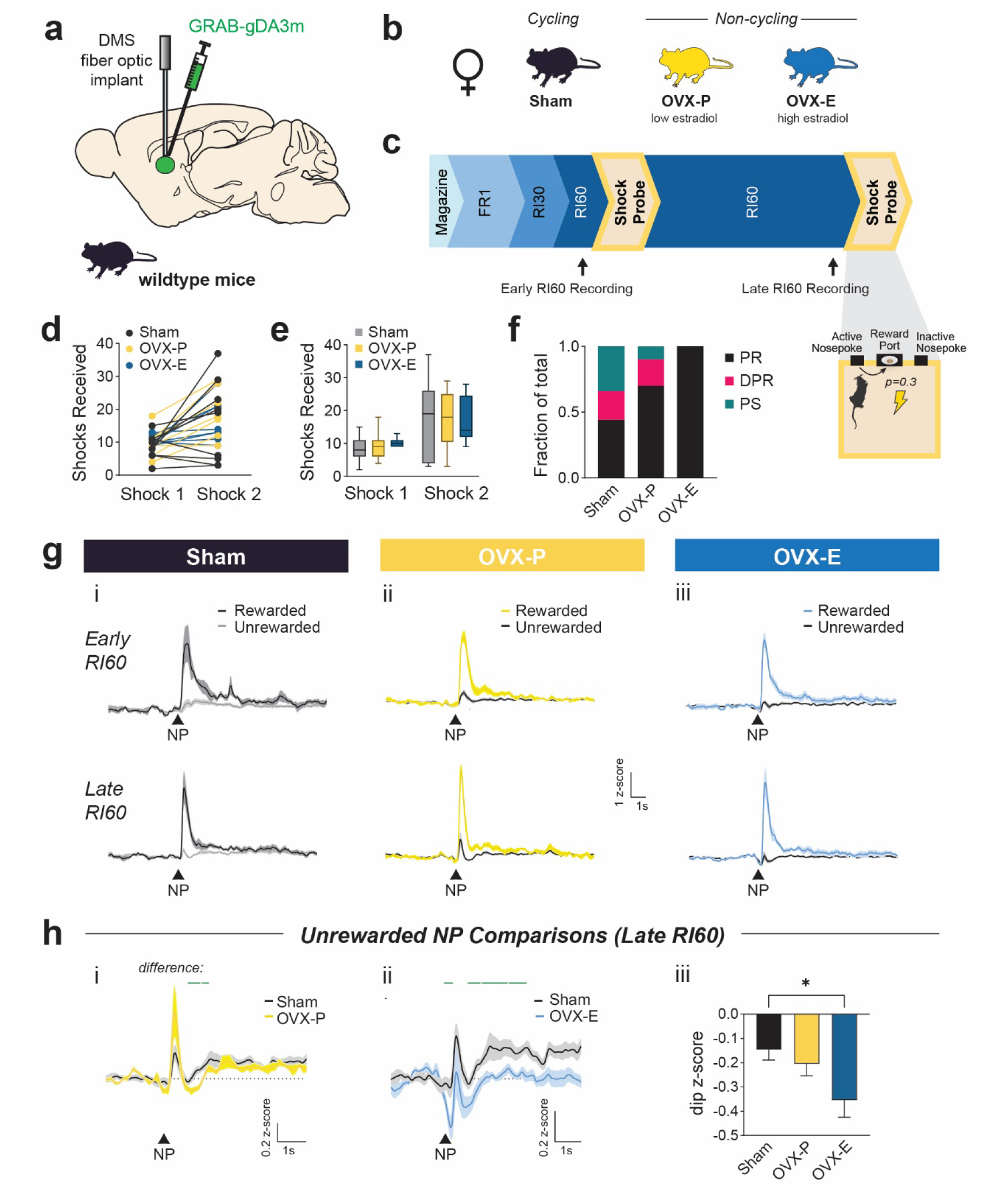
Chronic estradiol treatment increases vulnerability to punishment-resistant reward-seeking and amplifies DMS dopamine dips during unrewarded actions. A. Schematic of viral injection and probe placement strategy for recording DMS dopamine signals using GRAB-gDA3m fiber photometry in wildtype mice. B. Schematic of mouse experimental groups. Sham mice underwent a sham ovariectomy surgery and subcutaneous implantation of a placebo pellet. OVX-P mice underwent an ovariectomy and placebo pellet implantation. OVX-E mice underwent an ovariectomy and slow-release estradiol-17β pellet implantation. C. Schematic of the operant behavior training schedule. Shock probes introduced a risk of footshock for reward-seeking nosepokes (probability=0.3). Black arrows denote sessions early and late in RI60 training when fiber photometry signals were recorded. D-E. Shocks received for mice in Sham (black, n = 9), OVX-P (yellow, n=9) and OVX-E (blue, n=9) groups during the early (Shock 1) and late (Shock 2) probes. Spaghetti plots show lines connecting individual mice over time. Box plots show the median (center line), interquartile range (box), and whiskers representing the minimum and maximum values for each group. F. Fraction of mice classified as punishment resistant (PR, black; n=4 Sham, n=6 OVX-P, n=9 OVX-E), delayed punishment resistant (DPR, pink; n=2 Sham, n=2 OVX-P, n=0 OVX-E), punishment sensitive (PS, teal; n=3 Sham, n=1 OVX-P, n=0 OVX-E). Fisher’s Exact Test: Sham vs OVX-E, p=0.029. G. Peri-event time histograms (PETHs) showing DMS dopamine responses to rewarded and unrewarded nosepokes during early (top) and late (bottom) RI60 sessions. Sham recordings (i) are shown on the left (black/gray, n = 7), OVX-P (ii) middle (yellow/gray, n=6) and OVX-E (iii) right (blue/gray, n =7). H. PETHs comparing Sham (black, n = 7) vs OVX-P (yellow, n = 6; i) and Sham vs OVX-E (blue, n =7; ii) DMS dopamine responses to unrewarded nosepokes during late RI60. Green lines above the traces indicate periods during which waveform analysis with bootstrapped confidence intervals indicated significant differences. *iii*, Average z-score during unrewarded nosepokes in Sham, OVX-P, and OVX-E groups. Main effect of treatment: p=0.04 ;Tukey’s Multiple Comparisons: Sham vs OVX-E, *p=0.038.

Although behavior during RI60 training was similar, we found differences in punishment-resistant reward-seeking, particularly in OVX-E females (Fig. 1d-f). To compare phenotypes, we separated mice into three categories – punishment resistant (PR), delayed punishment resistant (DPR), and punishment sensitive (PS) – based on the number of shocks they tolerated on the early and late shock probes. Category criteria were based on our previous study^7^ but renormalized to the females in our Sham group (see Methods for details). Both OVX groups were more likely than Sham to be categorized as PR: OVX-P mice were majority PR or DPR (PR=70%, DPR=20%) while OVX-E mice were 100% PR (Fig. 1f). In addition, OVX-E mice were significantly different from Sham on the early shock probe (Fisher’s exact test: p = 0.0294) and showed less variance (Levene’s test: F = 4.7184, p = 0.0452; Fig. 1d-e). Overall, we concluded that a non-cycling state, particularly a non-cycling chronic high estradiol state, predisposes female mice towards developing punishment-resistant reward-seeking, without impacting the acquisition of RI60 behavior.

We next analyzed our fiber photometry recordings to determine if changes in DMS dopamine signals accompanied the behavioral changes. Dopamine signals during an early and late RI60 session were compared for all groups (Fig. 1g). Surprisingly, we found no significant differences in the dopamine peaks following a rewarded nosepoke in OVX-P or OVX-E compared to Sham (Fig. S2e). However, there was a significant difference in the OVX-P dopamine signal following unrewarded nosepokes, beginning ∼1s after the nosepoke (Fig. 1h_i_). In OVX-E mice, where the behavioral promotion of punishment resistance was stronger, there was also a significant difference following unrewarded nosepokes. The magnitude of the dopamine dip immediately following an unrewarded nosepoke was significantly greater (Fig. 1h_ii-iii_; Tukey’s multiple comparisons of peak dip z-score: p = 0.038), and there was an additional suppression of subsequent dopamine dynamics over the next several seconds (Fig. 1h_ii_). These data suggested that DMS dopamine dips on unrewarded nosepokes could be creating punishment-resistant reward-seeking. Such a role would be consistent with our prior work^7^; however, it was an untested causal hypothesis. Therefore, we set out to test it using optogenetics.

We used NpHR to inhibit DMS dopamine terminals during unrewarded nosepokes, hypothesizing that optogenetically creating a larger dopamine dip during unrewarded nosepokes would promote punishment-resistant reward-seeking over the course of RI60 training. We included both male and female mice in this experiment because it was unclear whether sex differences would be observed in the optogenetic effects. AAV5-EF1α-DIO-eNpHR3.0-EYFP (or fluorophore-only control virus AAV5-EF1α-DIO-EYFP) was injected bilaterally into the medial SNc of DAT-IRES-cre mice to preferentially target DMS-projecting dopamine neurons^22^. Fiber optic cannulas for light delivery were implanted bilaterally in the DMS to inhibit DMS dopamine terminals (Fig. 2a, S3a-b). A 1-second continuous light pulse (625nm) was administered during unrewarded nosepokes beginning with RI30 training (Fig. 2b). No optogenetic inhibition was delivered during the shock probes. To verify that our inhibition strategy amplified temporally precise dopamine dips without rebound effects, we combined fiber photometry and optogenetics to record DMS dopamine during optogenetic inhibition. For this test, we randomly interleaved unrewarded nosepokes on which inhibition was applied vs not applied within a single RI60 session. We saw that the NpHR inhibition effectively accentuates a dip in DMS dopamine during unrewarded nosepokes (Fig. 2c).

**Figure 2.**
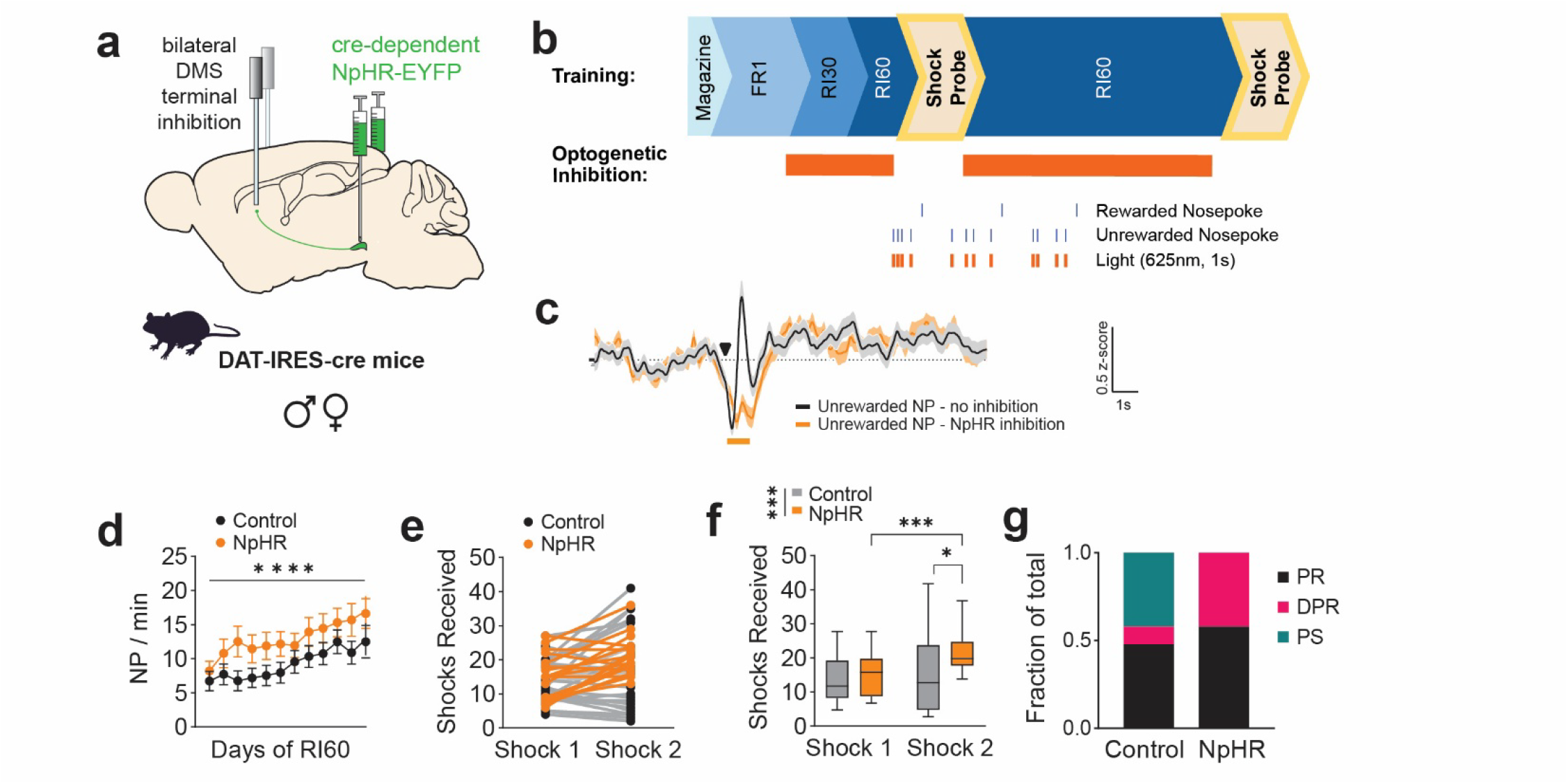
DMS Dopamine Terminal Inhibition on Unrewarded Nosepokes Promotes Punishment-Resistant Reward-Seeking. A. Schematic of the viral injection and probe placement strategy in DAT-IRES-cre mice for bilateral inhibition of DMS dopamine terminals using NpHR3.0. B. *Top*, Schematic of the operant behavior training schedule. Orange bars show sessions when optogenetic inhibition was applied. *Bottom*, schematic of the optogenetic inhibition applied following unrewarded nosepokes during RI60 training sessions. C. Peri-event time histogram (PETH) showing DMS dopamine (GRAB-gDA3m) recorded in response to unrewarded nosepokes during the RI60 task. For this test, trials with inhibition (orange) and no inhibition (black) were interleaved to validate the effects of inhibition. Black arrowhead indicates the time of unrewarded nosepoke. Orange bar indicates the time of NpHR inhibition on trials when it was delivered. D. Average nosepokes (NP) per minute across days of RI60 training for NpHR (orange, n=19) and control (black, n=29) mice. ****p = 0.0001 (main effect of day). E-F. Shocks received for mice in NpHR (orange, n=19) and control (black, n=29) groups. Spaghetti plots show lines connecting individual mice over time. Box plots show the median (center line), interquartile range (box), and whiskers representing the minimum and maximum values for each group. *p = 0.0131, ***p=0.0005. G. Fraction of mice classified as punishment resistant (PR, black; n=11 NpHR, n=14 control), delayed punishment resistant (DPR, pink; n=8 NpHR, n=3 control) and punishment sensitive (PS, teal; n=0 NpHR, n=12 control).

Both NpHR and control mice significantly increased their nosepoke rates across days of training but did not show differences in escalation (two-way mixed-effects ANOVA, F= 12.94, p<0.0001; Fig. 2d). NpHR mice performed significantly more port entries per minute compared to controls (Two-way mixed-effects ANOVA, F = 9.675, p = 0.043; Fig. S3c). However, there were no significant differences between the groups in rewards earned per session and nosepokes per reward (Fig. S3d-e). Importantly, there were no differences in the number of light stimulations received between the groups (Fig. S3f).

Turning to the shock probes, NpHR mice had a highly significant increase in the number of shocks received between Shock 1 and Shock 2 (Sidak’s multiple comparisons test: ***p = 0.0005; Fig. 2e-f). Further, we found a significant main effect of NpHR, and a significant increase in shocks received on Shock 2 in NpHR vs control mice (Two-way ANOVA main effect of virus: F=4.46, p=0.04; Sidak’s multiple comparisons test *p=0.0131) (Fig. 2e-f). All NpHR mice were categorized either as PR or DPR, while control mice still displayed a distributed range of phenotypes (Fig. 2g).

When we included sex as a factor in our analysis, we did not find a main effect of sex or sex by NpHR interaction (Three-way ANOVA: sex, p=0.08; sex by NpHR, p=0.06). Although this result suggests that optogenetic inhibition is effective in both sexes, we also analyzed male and female data separately given our interest in estradiol (Fig. S4). We observed a larger effect size in females (males η^2^p = 0.17, p=0.048; female η^2^p=0.38, p=0.0009). Taken together, our data confirm a causal link between DMS dopamine dips on unrewarded nosepokes and the development of punishment-resistant behavior, meaning this is a plausible explanatory factor for the effects of OVX-E.

In sum, our observation of hormone-dependent differences in DMS dopamine dips on unrewarded nosepokes in female mice led us to a generalizable conclusion for males and females, highlighting an important causal behavioral role for dopamine dips. Contrary to the predictions of reward prediction error theory, dopamine dips did not decrease performance of the action (nosepoking) but instead changed the degree of learning over time about the value of the action. Our finding that *decreases* in dopamine promoted persistence in reward-seeking in the face of punishment thus challenges prevailing theory, but it is consistent with proposals that dopamine controls policy learning or operates through an adjusted net contingency algorithm^23–25^. In short, DMS dopamine here seems to suppress learning about action value rather than reducing estimates of action value directly.

Our finding that DMS dopamine dips can be naturally accentuated under chronic estradiol treatment (OVX-E) also addresses a translationally relevant gap in our understanding of the role of hormonal cycling in calibrating female dopamine function. Many women around the world use hormonal birth control that disrupts cyclic changes in estrogen, and women in perimenopause and menopause face changes due to low estrogen and decisions about whether to take hormone replacements. Despite the known impact of hormonal treatments on mood and psychiatric vulnerability, these populations of women are not well modeled in animal studies that consider only young cycling females, and there is still little known about how the lack of cycling affects dopamine signaling ^26,27^. Our finding is consistent with work indicating that estradiol modulates dopamine transporter (DAT) function ^19,20,28,29^, since clearance rates of dopamine will impact the magnitude of dips during pauses in firing. The current observation thus makes a strong case for further studies of how DAT function is regulated under different cycling and non-cycling hormonal states, which could influence women’s vulnerability to psychiatric disorders.

### METHODS

#### Experiment Model and Subject Details

Adult (10+ weeks) female and male mice were group housed by sex on a reverse 12:12h light/dark cycle with ad libitum access to food and water prior to operant training. Female WT (C57BL/6J) mice obtained from The Jackson Laboratory were used in the ovariectomy experiments.

Cagemates were randomly assigned to experimental groups (ovariectomy and estradiol pellet - 9; ovariectomy and placebo pellet - 9 ; sham and placebo pellet - 9). Heterozygote transgenic DAT-IRES-Cre mice were used for NpHR optogenetic experiments and obtained by crossing (DAT)::IRES-Cre knockin mice (JAX006660) and WT (C57BL/6J) mice. Littermates were randomly assigned to experimental groups (NpHR - 19, EYFP control - 29). All experiments were approved by the Northwestern University Institutional Animal Care and Use Committee.

### Methods Details

#### Ovariectomy Surgery

Ovariectomy surgery took place under isoflurane anesthesia (Henry Schein). Mice were anesthetized in an isoflurane induction chamber at 3-4% isoflurane then placed upside down, so the abdomen was exposed. Anesthesia was maintained at 2-3% isoflurane. Mice were injected with meloxicam (Covetrus, 20 mg/kg) and buprenorphine (Ethiqa XR, 3.25mg/kg) before surgery began. Hair on the abdomen was removed with Nair and skin was disinfected with alcohol and povidone-iodine. A small incision was made in the middle of the abdomen, cutting skin and muscle. Sterile forceps were used to locate the ovary and a dissolvable suture (Ethicon, 7-0 coated vicyrl 18”, J488G) was used to secure the distal part of the uterine horn to prevent bleeding. The ovary was resected, and the uterine horn was placed back in the abdominal cavity. The same procedure was repeated to remove the second ovary. The muscle was sutured with absorbable sutures, and the skin was closed with non-absorbable surgical sutures (Ethicon,6-0 Ethilon 18”, 697H). After surgery, mice were allowed to recover until ambulatory on a heated pad, then returned to their homecage with moistened chow or DietGel available. The mice were checked after 24 hours.

#### Stereotaxic Surgery

Fiber optic implants and viral infusions performed on adult mice took place under isoflurane anesthesia (Henry Schein). Mice were anesthetized in an isoflurane induction chamber at 3-4% isoflurane, then placed on a stereotaxic frame (Stoetling), where anesthesia was maintained at 1-2% isoflurane. Mice were then injected with meloxicam (Covetrus, 20 mg/kg) and, for OVX experiments, with long-acting buprenorphine (Ethiqa XR, 3.25mg/kg) before surgery began. For OVX experiments, stereotaxic surgery took place one week after ovariectomy. Nair was used to remove hair from the incision site, and alcohol and povidone-iodine were used to disinfect exposed skin. Prior to incision, bupivacaine (Hospira, 2 mg/kg) was injected subcutaneously at the incision site. A sterile scalpel was used to open the scalp, and holes were drilled in the skull at appropriate stereotaxic coordinates. Viruses were infused at 100 nL/min through a blunt 33-gauge injection needle using a syringe pump (World Precision Instruments). The needle was left in place for 5 min following the end of the injection, then slowly retracted to avoid leakage up the injection tract. Implants were secured to the skull with Metabond (Parkell) and Flow-it ALC blue light-curing dental epoxy (Pentron).

After surgery, mice were allowed to recover until ambulatory on a heated pad, then returned to their homecage with moistened chow or DietGel available. The mice were checked after 24 hours and provided with another dose of meloxicam. Mice then recovered for two weeks before behavioral experiments began.

#### Subcutaneous Pellet Implant Surgery

Pellets were implanted during stereotaxic surgeries. Nair was used to remove hair from the neck. Alcohol and povidone-iodine were used to disinfect the skin. A small incision was made and a Placebo or slow-release Estradiol-17β (0.01mg, 90-day release, Innovative Research of America, Cat. No. NE-121) was implanted subcutaneously in the space between the shoulder and neck. The incision was closed using non-absorbable sutures.

#### Operant Behavior

For the duration of operant training, mice were food-restricted to 85% of their ad libitum body weight. Mice were trained to retrieve sugar pellet rewards (20mg, Sucrose, Dustless Precision Pellets, Bio-Serv) from a magazine port. During magazine training, pellets were delivered to the port on a random interval (RI60) schedule non-contingently for one hour. Mice were then trained to associate nosepoking with reward on a fixed ratio (FR1) schedule where both nosepokes delivered a reward. Mice had to retrieve the reward (as measured by making a port entry following a rewarded nosepoke) before they could earn the next reward. After mice received >25 rewards on one side they were trained on FR1 on their preferred side only, with nosepokes on the other side having no consequence. When mice received >30 rewards for a minimum of two consecutive days on the preferred side they moved to a random interval 30 (RI30) schedule of reinforcement. Two female control mice that did not reach FR1 criterion after 14 days of training were removed from the study. Mice were trained on RI30 until they earned >30 rewards in one hour, then trained on RI60. All operant training sessions lasted one hour or until 50 rewards were earned.

#### Shock Probes

To evaluate punishment-resistant reward-seeking, mice were subjected to footshock probes at both early and late stages of RI60 training. During the shock probes, which took place under an FR1 schedule of reinforcement, a mild footshock (0.2mA, 1s) was paired with a subset of rewarded nosepokes on a RR3 schedule, so that, on average, every third rewarded nosepoke was paired with a footshock. The first five rewarded nosepokes were never paired with shock. During shock probes, the session ended after 60 minutes, or if a mouse did not poke on the rewarded side for >10 minutes. There was no maximum number of rewards.

#### Optogenetics

For inhibitory optogenetics experiments, mice received 1 μl per side of AAV5-EF1α-DIO-eNpHR3.0-EYFP (1.1e13 GC/mL, Addgene, Lot:v32533) or the control fluorophore-only virus AAV5-EF1α-DIO-EYFP (3.5e12 GC/mL, UNC Vector Core, Lot: AV4310K) in bilateral medial SNc (AP −3.1, ML 0.8, DV −4.7).

Bilateral fiber optic implants (Prizmatix; 500μm core, 0.66 NA) were placed in DMS (AP 0.8, ML ±1.5, DV −2.8). Starting during RI30 training and continuing until the end of operant behavior, unrewarded nosepokes were paired with a continuous pulse of orange/red light (625nm, 1s, 15 mW) generated by an LED light source and pulse generator (Prizmatix). After each instance of light delivery, there was a 6s timeout period in which subsequent unrewarded nosepokes would not be paired with light stimulation to prevent excessive light exposure. Light was not delivered during shock probes.

#### Fiber Photometry

For fiber photometry experiments, mice received a 500 nL infusion of AAV1-hsyn-GRAB-gDA3m (1.3e13 GC/mL, Addgene, 208698) in the DMS (AP 0.8, ML ±1.5, DV −2.8) in one hemisphere, with the hemisphere counterbalanced between mice. Fiber optic implants (Doric Lenses; 400um; 0.48NA) were implanted in the DMS at the same coordinates. Dopamine signals were recorded on the first and last days of RI60 training. All recordings were performed using a fiber photometry rig with optical components from Doric lenses and controlled by real-time processors from Tucker Davis Technologies (TDT). Behavioral events from Med Associates were fed into the real-time processor as TTL signals for alignment with neural data. GuPPy, an open-source Python-based photometry data analysis pipeline ^30^, was used to process fiber photometry data and align signals to specific time-locked events.

#### Perfusions and Histology

After the conclusion of operant behavior, euthasol (Virbac, 1mg/kg) was administered to mice i.p. to induce rapid onset unconsciousness and death. Once unresponsive to a firm toe pinch, an incision was made in the abdomen to open the middle body cavity and the heart was exposed. An injection needle connected to a peristatic pump (Mini Pump Variable Flow; Fisher Scientific) via tubing was inserted into the left ventricle of the heart, the right atrium was punctured, and PBS followed by 4% PFA was infused into the left ventricle as the mouse was exsanguinated. The mouse was decapitated, and the brain was removed and fixed in 4% PFA overnight at 4 °C. Brains were then transferred to a solution of 30% sucrose in PBS (w/v). Tissue was sectioned on a freezing microtome (Leica) at 50 μm, stored in cryoprotectant (30% sucrose, 30% ethylene glycol, 1% polyvinyl pyrrolidone in PBS) at 4 °C until immunostaining. Anti-GFP staining was performed on free-floating sections to amplify signals from GRAB-gDA3m and eNpHR3.0-EYFP. Sections were blocked in 3% normal goat serum in PBS for 1 h at room temperature. Primary antibody staining was performed using 1:500 Rabbit anti-GFP primary antibody (Invitrogen, A11122) in blocking solution at 4°C for 48 hrs. Secondary staining was performed using 1:1000 donkey anti-rabbit Alexa Fluor 488 (Jackson ImmunoResearch) in blocking solution at room temperature for 2 hrs. Tissue sections were mounted onto slides in PBS, coverslipped (Fisherbrand, Cat. No. 12-550-05) with DAPI Fluoromount-G (SouthernBiotech).Slides were imaged using a fluorescent microscope (Keyence BZ-X710) equipped with 5x, 10x, and 40x air-immersion objectives. Probe placements were determined by comparing their location to the Allen Mouse Brain Atlas.

#### Vaginal Swabbing and Estrous State Analysis

After the conclusion of operant behavior and before sacrifice, vaginal lavage was performed to confirm the effects of estradiol treatment. A latex bulb was placed on the end of a sterile 200um tip and approximately 100-120ul of Milli-Q water was used to fill the sterile tip. The Milli-Q water was placed at the opening of the vaginal canal of the mice and then withdrawn back into the tip and this action was repeated 3-4 times to ensure cell collection. The resulting fluid was placed onto a glass slide (Fisherbrand,Cat. No. 12-550-05) and the smear was allowed to dry at room temperature. Glycerol (Fisher Bioreagents, Cat. No. BP229-1) and a coverslip (Fisherbrand, Cat. No. 125-410-55) were applied and left to dry overnight. Brightfield images were taken at 40x magnification (Keyence BZ-X710) and used to identify estrous state. Proestrus is categorized by the predominance of nucleated epithelial cells while diestrus is categorized by the predominance of leukocytes. In estrus there is the presence of cornified squamous epithelial cells packed together and Metestrus included both leukocytes and cornified squamous epithelial cells ^31^. OVX-E mice were excluded if the images from vaginal lavage resembled metestrus or diestrus, indicating that the estradiol pellet had not been effective at creating a high estrous state.

### Quantification and Statistical Analysis

#### Behavioral analysis

Behavioral data (e.g. nosepokes, port entries) were collected automatically by MED-PC software (Med Associates). After RI60 training, mice were sorted into punishment resistant (PR), delayed punishment resistant (DPR) and punishment sensitive (PS) groups based on post hoc analysis of their performance during the shock probes. Mice were sorted by a median split on the number of shocks received by Sham (OVX experiments) or Control (NpHR experiments) mice on shock 1. We separated mice into PR (receiving >median shocks on both the early and late shock probes), DPR (receiving <median shocks in the early probe but increasing >85% on the late shock probe, to >median), or PS (receiving <median shocks on the early and late shock probes).

#### Fiber photometry analysis

GuPPy ^30^ was used to analyze fiber photometry signals time-locked to specific behavioral events using default settings. In brief, raw data were passed through a zero-phase digital filter and a least-squares linear fit was applied to the 405nm control signal to align it to the 465nm signal. The following formula was used to calculate ΔF/F: (signal - fitted control) /(fitted control). Z-scores were calculated by subtracting the mean ΔF/F calculated across the entire session and dividing by the standard deviation.

Peristimulus time-histogram (PSTH) parameters were calculated from -10 to 20 seconds and baseline subtraction was performed using a baseline period from -5 to 0 seconds. H5 output files were converted to csv files and values were entered into GraphPad Prism to visualize traces. Bootstrapped confidence interval waveform analysis (using 95% confidence intervals) was conducted on photometry traces downsampled to 100 Hz to identify significant deviations between fiber photometry signals of Sham, OVX-E and OVX-P mice.

Bootstrapped confidence interval identified significant deviations only when the bounds of 95% CI did not include zero and did not overlap for more than 20 consecutive data points (200 ms) ^32^.

#### Other statistical methods

One-way, two-way and three-way ANOVAs and multiple comparison analyses were performed using Prism 11 software. Sidak’s and Tukey’s multiple comparison analyses were performed only when statistically significant main effects were found. Variance was verified by Levene’s test and categorical association was evaluated using Fisher’s exact test and both tests were performed using custom-written code (Python, version 3.9.19).

## ACKNOWLEDGEMENTS

We thank the Lerner laboratory for helpful discussions and critical feedback throughout the project, Hayden Sikora for his early efforts towards establishing this research direction under the supervision of J.L.S, Louis Van Camp and Briana Hryhorysak for supporting the laboratory’s general operations, Venus Sherathiya for technical support using GuPPy for fiber photometry analysis, and the Center for Comparative Medicine at Northwestern University for providing animal care and husbandry.

## AUTHOR CONTRIBUTIONS

N.M., J.L.S. and T.N.L. conceived the studies. N.M., J.L.S., and T.S. conducted the experiments. N.M. and J.A.N. analyzed the data. N.M. and T.N.L. wrote the manuscript. T.N.L. provided oversight and support for the project.

## DATA AVAILABILITY

Fiber photometry, behavioral, and optogenetic data supporting the findings of this study will be deposited in the DANDI Archive prior to publication and will be made available at [link TBD]. In the interim, data are available from the corresponding author upon reasonable request.

## CODE AVAILABILITY

No custom code was used in this study. Data were analyzed using GuPPy (https://github.com/LernerLab/GuPPy) and GraphPad Prism.

**Figure S1.**
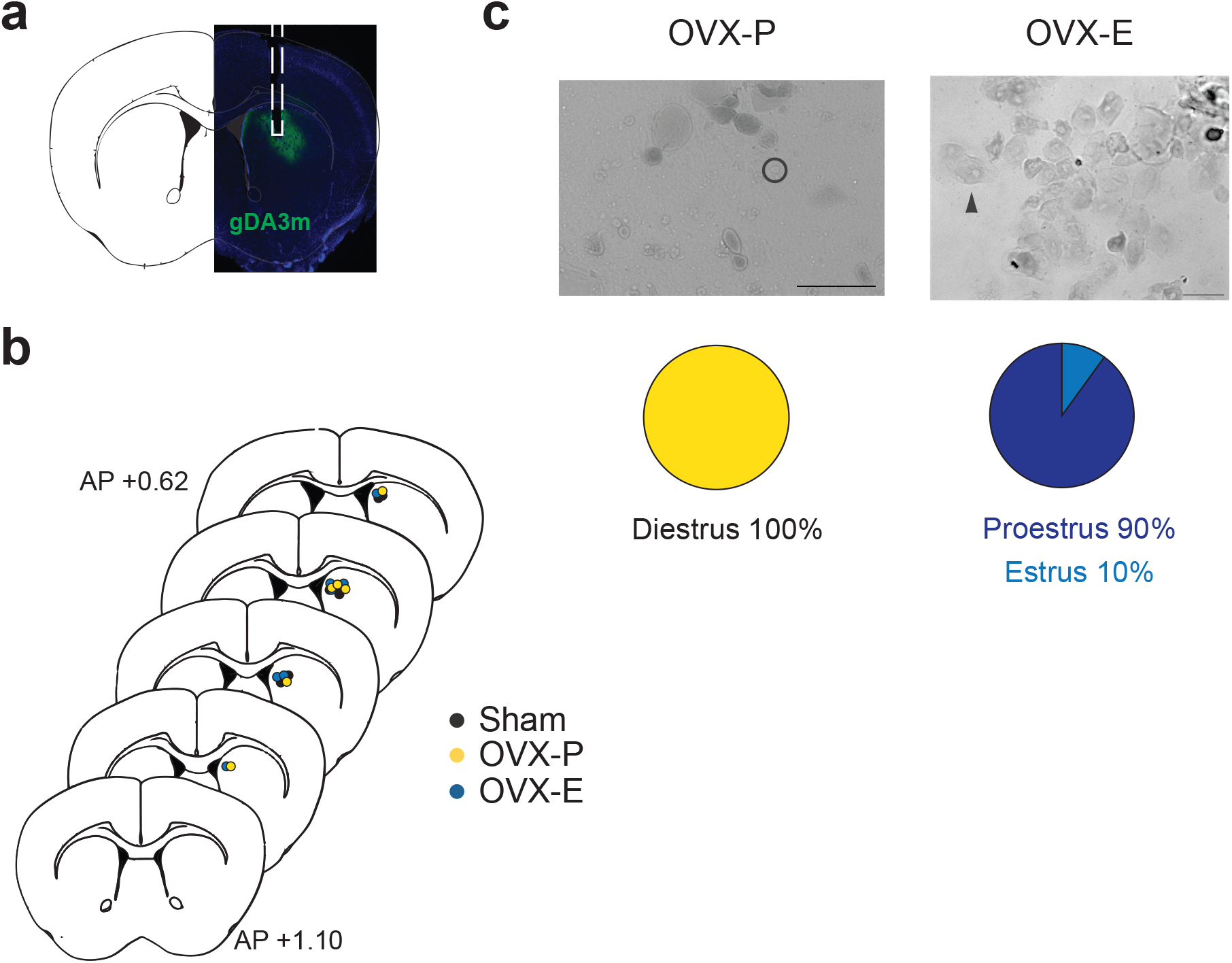
Estrous Tracking, Viral Expression, and Probe Placements for Estrous State Experiments (Related to Figure 2) A. Representation 10x image of the striatum showing GRAB-DA3m expression (green) and probe placement (white dashed line). B. Probe placements for all mice in Figure 1 (Sham: black, OVX-P: yellow, OVX-E: blue). C. Representative 40x images of cells (leukocytes (circle) and nucleated epithelial (black arrow)) from vaginal lavage of OVX-P (right) and OVX-E (left) mice to determine estrous state. The pie charts below show the percentages of mice classified as diestrus, proestrus, or estrus in both groups.

**Figure S2.**
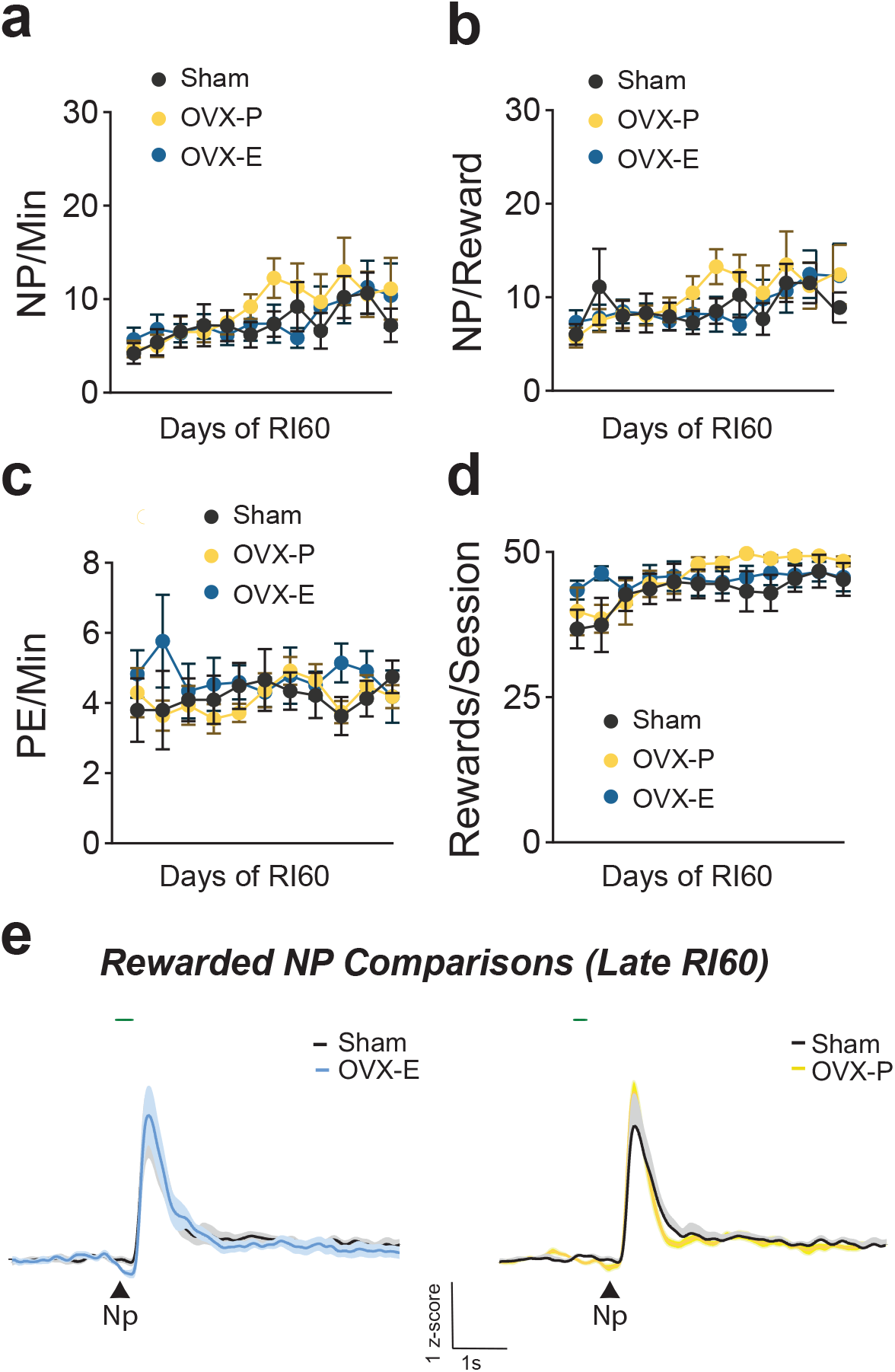
Additional Effects of Estrous State Manipulations (Related to Figure 1) A. Average nosepokes (NP) per minute for Sham (black, n = 9), OVX-P (yellow, n=9) and OVX-E (blue, n=9) across days of RI60 training. B. Average nosepokes (NP) per reward for Sham (black, n = 9), OVX-P (yellow, n=9) and OVX-E (blue, n=9) across days of RI60 training. C. Average port entries (PE) per minute for Sham (black, n = 9), OVX-P (yellow, n=9) and OVX-E (blue, n=9) across days of RI60 training. D. Average rewards per session for Sham (black, n = 9), OVX-P (yellow, n=9) and OVX-E (blue, n=9) across days of RI60 training. E. PSTHs comparing Sham (black, n = 7) vs OVX-E (blue, n=7, right) and Sham vs OVX-P (yellow, n=6, left) DMS dopamine responses to rewarded nosepokes during late RI60. Green lines above the traces indicate periods during which waveform analysis with bootstrapped confidence intervals indicated significant differences.

**Figure S3.**
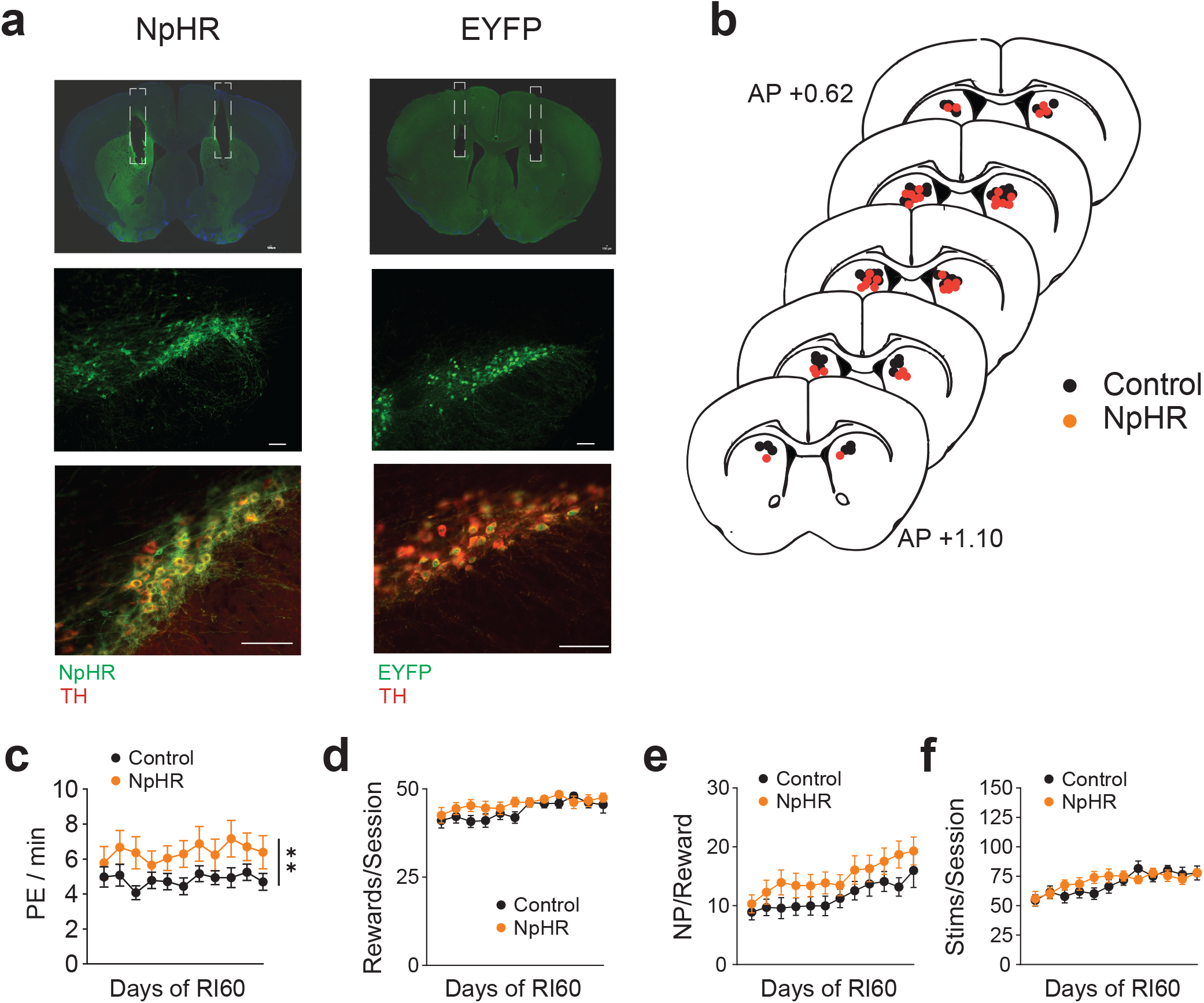
Viral Expression, Probe Placements, and Additional Behavioral Results for DMS Dopamine Terminal Inhibition Experiments (Related to Figure 3) A. Representative 4x images of probe placement in the DMS for NpHR (left) and Control (right) mice. Representative 10x images showing expression of NpHR3.0 and TH (left) or EYFP (right) in substantia nigra pars compacta (SNc) dopamine neurons. Scale bar for all images is 100um. B. Probe placements for all mice in Figure 2 (Control: black, NpHR: orange). C. Average port entry (PE) per minute across days of RI60 training for NpHR (orange, n=19) and control (black, n=29) mice. **p = 0.004 (main effect of treatment). D. Average rewards per session for Control (black, n= 29) and NpHR (orange, n =19) mice across days of RI60 training D. Average nosepokes (NP) per reward for Control (black, n= 29) and NpHR (orange, n =19) mice across days of RI60 training. E. Average number of light stimulations per session for Control (black, n= 29) and NpHR (orange, n =19) across days of RI60 training.

**Figure S4.**
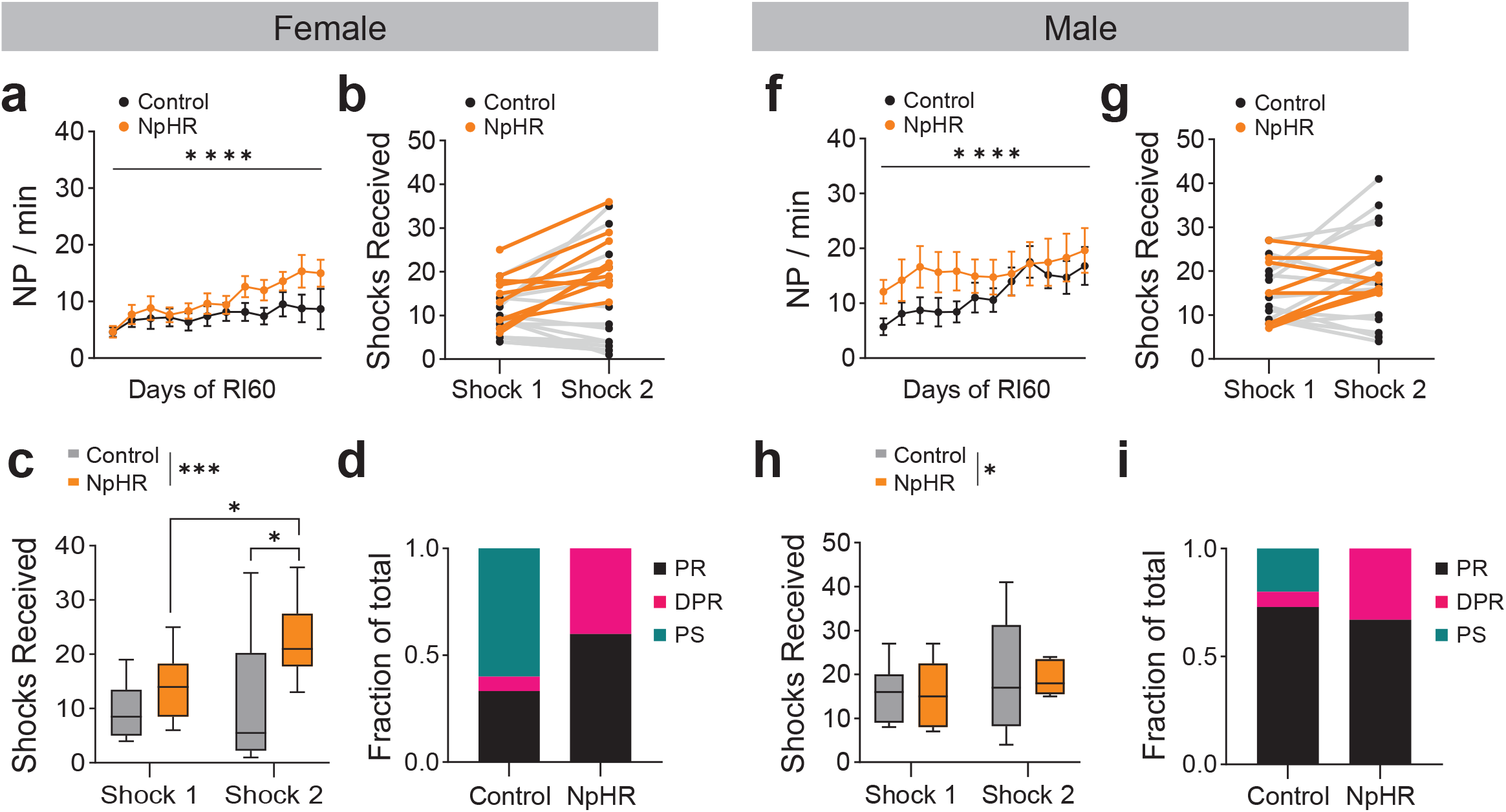
Effects of DMS Dopamine Terminal Inhibition by Sex (Related to Figure 3). A,F. Average nosepokes (NP) per minute for females (A; Control n=15, NpHR n=10) and males (F, Control n=14, NpHR n=9) across days of RI60 training. B-C,G-H. Shocks received for females (B-C; Control n=15, NpHR n=10) and males (G-H, Control n=14, NpHR n=9). Spaghetti plots show lines connecting individual mice over time. Box plots show the median (center line), interquartile range (box), and whiskers representing the minimum and maximum values for each group. A 3-way ANOVA indicated a main effect of time, but no main effect of sex for NpHR. Sidak’s multiple comparisons: NpHR females shock 1 vs shock 2, *p = 0.030; Control females shock 2 vs NpHR females shock 2, *p = 0.026). D,I. Fraction of female (D) and male (I) mice classified as punishment resistant (PR, black; Control female n=5, NpHR female n=6, Control male n=11, NpHR male n=6), delayed punishment resistant (DPR, pink; Control female n=1, NpHR female n=4, Control male n=1, NpHR male n=3) and punishment sensitive (PS, teal; Control female n=9, NpHR female n=0, Control male n=3, NpHR male n=0).

